# The FAAH inhibitor URB597 reduces cocaine seeking during conditioned punishment and withdrawal

**DOI:** 10.1101/2023.03.13.532386

**Authors:** Laia Alegre-Zurano, Alba García-Baos, Adriana Castro-Zavala, Ana Martín-Sánchez, Mireia Medrano, Ines Gallego-Landin, Olga Valverde

## Abstract

The endocannabinoid system is prominently implicated in the control of cocaine reinforcement due to its relevant role in synaptic plasticity and neurotransmitter modulation in the mesocorticolimbic system. The inhibition of fatty acid amide hydrolase (FAAH), and the resulting increase in anandamide and other N-acylethanolamines, represents a promising strategy for reducing drug seeking. In the present study, we aimed to assess the effects of the FAAH inhibitor URB597 (1 mg/kg) on crucial features of cocaine addictive-like behaviour in mice. Therefore, we tested the effects of URB597 on acquisition of cocaine (0.6 mg/kg/inf) self-administration, compulsive-like cocaine intake and cue-induced drug-seeking behaviour during withdrawal. URB597 reduced cocaine intake under conditioned punishment while having no impact on acquisition. This result was associated to increased cannabinoid receptor 1 gene expression in the ventral striatum and medium spiny neurons activation in the nucleus accumbens shell. Moreover, URB597 reduced cue-induced drug-seeking behaviour during prolonged abstinence and prevented the withdrawal-induced increase in FAAH gene expression in the ventral striatum. In this case, URB597 decreased activation of medium spiny neurons in the nucleus accumbens core. Our findings evidence the prominent role of endocannabinoids in the development of cocaine addictive-like behaviours and support the potential of FAAH inhibition as a therapeutical target for the treatment of cocaine addiction.

## INTRODUCTION

The transition from recreational cocaine use to pathological abuse is characterized by loss of control over cocaine intake, compulsive cocaine consumption despite negative consequences, abstinence-induced negative states, and increased reactivity to drug-associated cues that leads to relapse [1–3]. Although preclinical and clinical research during the last decades have increased our understanding about different traits of cocaine addiction and their neurobiological bases, many matters remain unsolved.

Given its involvement in a wide range of neural processes, including synaptic plasticity and neurotransmitter release [4], the endocannabinoid system (ECS) holds great potential as a modulator of addiction-related processes such as mood, learning and reinforcement [5,6]. The ECS primarily consists of receptors (cannabinoid receptor 1 [CB1R] and 2 [CB2R]), endogenous ligands (anandamide [AEA] and 2-arachidonoylglycerol [2-AG]) and the enzymes responsible for their synthesis (N-acylphosphatidylethanolamine-specific phospholipase D [NAPE-PLD] and diacylglycerol lipase [DAGL]), and degradation (fatty acid amide hydrolase [FAAH] and monoacylglycerol lipase [MAGL]) [7]. The ECS controls the neuroplastic mechanisms that regulate the synapses from distal glutamatergic projections, primarily the medial prefrontal cortex (mPFC) and the amygdala, into medium spiny neurons (MSNs) within the nucleus accumbens (NAc). This occurs through endocannabinoid-mediated long-term depression (eCB-LTD), a mechanism lost after cocaine exposure [8–10]. Cocaine-induced alterations in the NAc are especially relevant since the NAc core and shell differentially modulate approach [11–13] and aversion [14,15] behaviours as well as their conditioned responses [16,17].

The inhibition of FAAH has recently been proposed as a potential treatment for drug addiction since endocannabinoids regulate excitatory and inhibitory neurotransmitter release in the mesocorticolimbic dopaminergic system, including the NAc. The FAAH inhibitor URB597 increases the endogenous levels of AEA as well as other N-acylethanolamines [18,19]. Such increase leads to the activation of the two canonical cannabinoid receptors CB1R and CBR2, but also other receptors belonging to the so-called expanded ECS, including transient receptor potential vanilloid 1 (TRPV1), G protein-coupled receptor 55 (GPR55) [7] and peroxisome proliferator-activated receptors (PPARs) [20]. Besides, URB597 has been reported to reduce tyrosine hydroxylase (TH) expression *in vitro* through CB1R- and FAAH-independent mechanisms [21]. Due to its pharmacological profile, URB597 does not elicit the typical spectrum of cannabinoid responses produced by direct CB1R agonism [22], but instead it exerts analgesic [23–25], anxiolytic [26–30] and antidepressant-like effects [29,31–33]. Additionally, despite of some degree of controversy [34], most data suggests that URB597 has positive effects on cognition [35–39].

During the last decade, preclinical studies have provided evidence on the protective effects of FAAH inhibition against nicotine [40–44], alcohol [38,45–48] and morphine [49] abuse. However, the role of URB597 as a potential treatment for cocaine use disorder remains to be fully addressed. URB597 does not alter cocaine-induced hyperlocomotion [50], but it does protect against cocaine-induced neurotoxicity [51]. Moreover, URB597 increases the preference for the cocaine-paired compartment in the conditioned place preference (CPP) paradigm [52], whereas in the intravenous self-administration model, it elicits different effects depending on the administration protocol. Therefore, acute URB597 treatment does not affect cocaine self-administration in rats [53] or squirrel monkeys [54], but it decreases both cue- and cocaine-induced reinstatement in rats [53]. Besides, chronic URB597 treatment during abstinence improves extinction and ameliorates cue- and stress-induced reinstatement in rats [55]. Additionally, URB597 efficiently prevents cocaine-induced depression of MSNs excitability in the NAc shell, but not in the ventral tegmental area (VTA) [44].

Despite the encouraging evidence about the therapeutic potential of FAAH inhibition in addressing cocaine-related behaviours, a thorough analysis is still needed to determine its effects of on core features of cocaine addiction, such as compulsivity and craving, as well as its impact on neuronal activation. In this study, we addressed these issues by evaluating the effects of URB597 on cocaine consumption under the risk of punishment and cue-induced cocaine-seeking behaviour during withdrawal. Additionally, we investigated the effects of URB597 on neuronal activation in the NAc as well as gene expression levels of the main components that participate in the endocannabinoid-mediated plasticity.

## MATERIALS AND METHODS

### Animals

Male CD1 mice (postnatal day 56) were purchased from Charles River (Barcelona, Spain) and maintained in the animal facility (UBIOMEX, PRBB) in an inversed 12-hour light-dark cycle (lights on 19:30-7:30). Mice were housed at a stable temperature (22 °C ± 2) and humidity (55% ± 10%), with food and water *ad libitum*. They were allowed to acclimatize to the new environmental conditions for at least four days prior to surgery. Animal care and experimental protocols were approved by the Animal Ethics Committee (CEEA-PRBB), following the European Community Council guidelines (2016/63/EU).

### Materials

Cocaine HCl (0.6 mg/kg/inf) was purchased in Alcaliber S.A. (Madrid, Spain), and was dissolved in 0.9% NaCl. URB597 (1 mg/kg i.p.) was purchased in Merck Life Science (Madrid, Spain) and was dissolved in a vehicle solution composed by ethanol, cremophor E.L. (Sigma-Aldrich) and 0.9% NaCl (1:1:18). The URB597 dose was established based on previous studies [53].

### Surgery and cocaine self-administration

The surgical procedure was conducted as previously described [56,57] with minor modifications. The cocaine operant self-administration paradigm was adapted from previous studies [57,58] (see Supplementary material). Briefly, mice were trained to nosepoke to obtain cocaine. Nosepokes in the active hole resulted in the delivery of a 20 μL cocaine infusion over 2 seconds (0.6 mg/kg/inf) and a stimulus light for 4 seconds, followed by a 15-second time-out period.

### Behavioural testing

#### Experiment 1: Effects of URB597on cocaine self-administration

Mice were trained to self-administer cocaine under a fixed ratio (FR) 1 schedule of reinforcement for 10 days (ACQ1-10). Mice were injected with either vehicle or URB597 30 minutes before every acquisition session (Fig. 1A). No performance-based exclusion criteria were applied.

**Fig 1.**
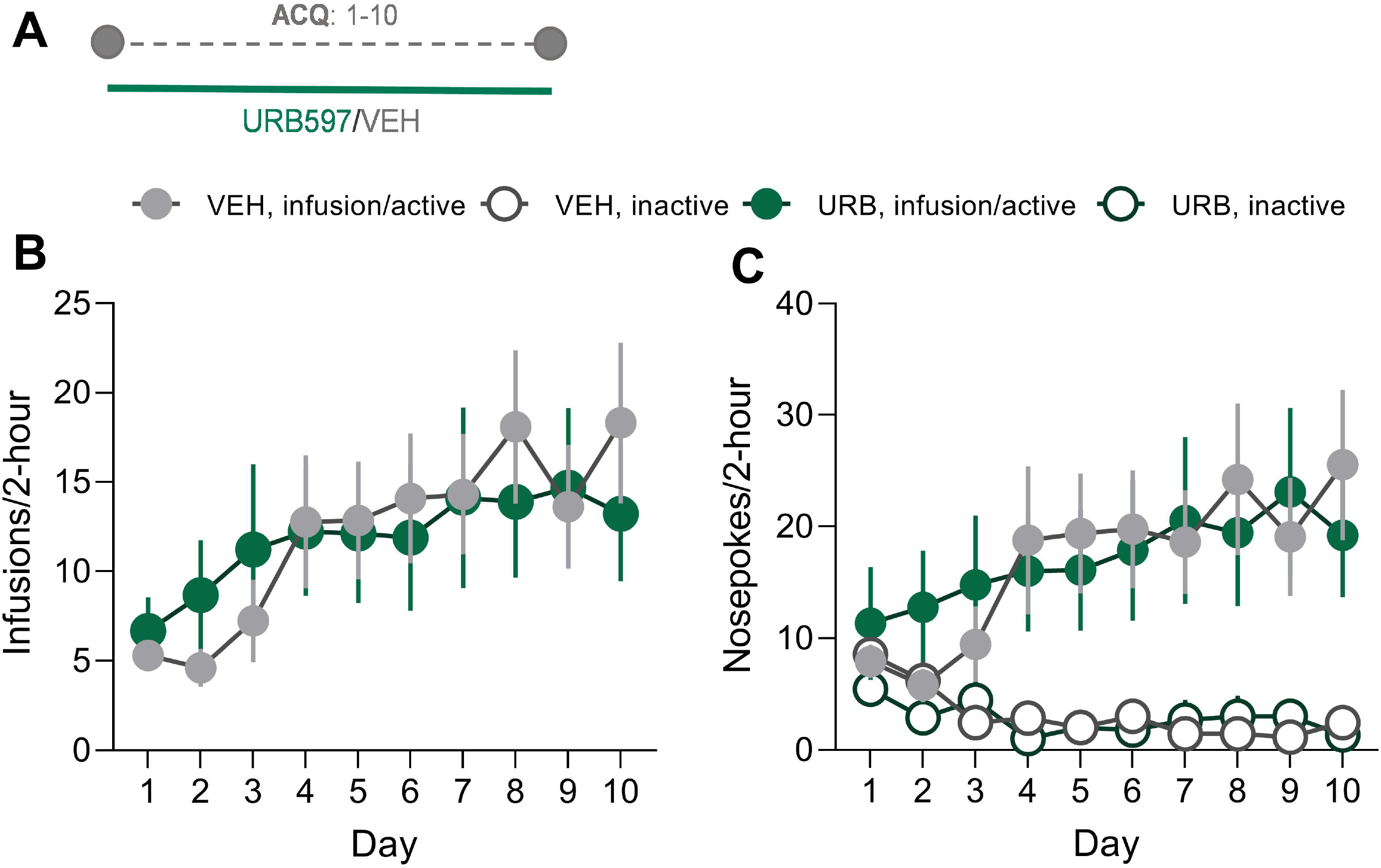
URB597 (1 mg/kg) does not modulate acquisition of cocaine self-administration. **A** Schematic timeline of the experiment 1. **B** Number of infusions and **C** nosepokes during the 10, 2-hour self-administration sessions (VEH n=13, URB n=9). Data are shown as mean ± SEM. ACQ: acquisition of cocaine self-administration, VEH: vehicle, URB: URB597.

#### Experiment 2: Effects of URB597 on cocaine self-administration under conditioned punishment

Mice underwent a 10-day acquisition of cocaine self-administration (ACQ1-10) under FR1 (Fig. 2A) and those who did not meet the following criteria on three consecutive days were excluded from the experiment: a minimum of 5 infusions and 65% of responses received in the active nosepoke. The acquisition phase was followed by three, 1-hour daily punishment sessions (PNS11-13), in which one third of the infusions were punished with a foot shock delivery (0.4 mA, 500 ms). The intensity of the foot shock was established based on experimental evidence (Supplementary Fig. 1). Additionally, a predictive stimulus consisting of the house light was presented immediately after the infusion preceding the foot shock and stayed on until the animal performed the active nosepoke leading to the punished infusion (Fig. 2B).

**Fig 2.**
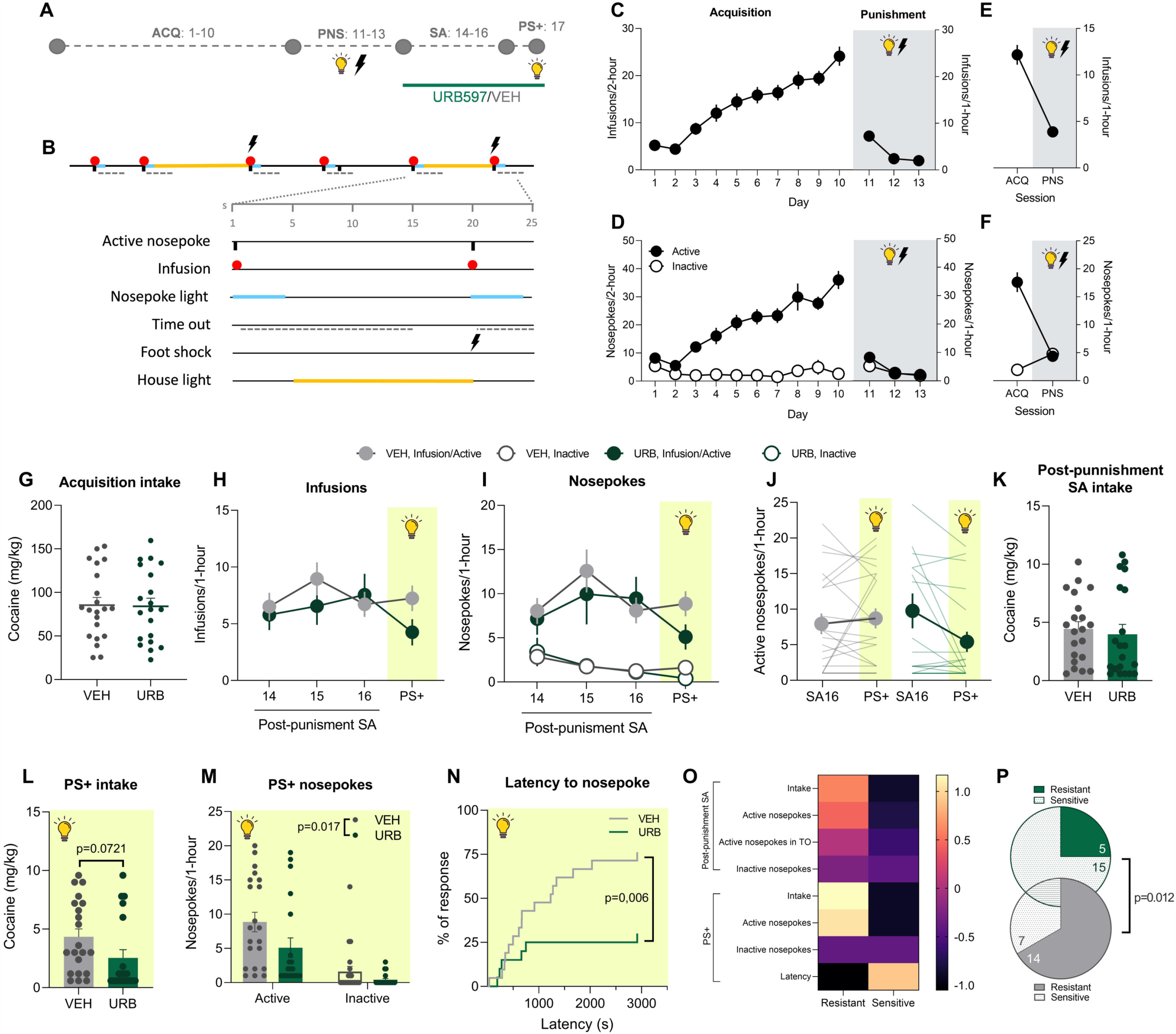
URB597 (1 mg/kg) decreases cocaine seeking when a punishment-predictive cue is present. **A** Schematic timeline of the experiment 2. **B** Schematic representation of the punishment protocol. **C** Number of infusions during the 10 2-hour acquisition sessions and the 3, 1-hour punishment sessions. **D** Mean of the number of infusions during the first hour of the last 3 acquisition sessions and the 3 punishment sessions. **E** Number of active and inactive nosepokes during the 10, 2-hour acquisition sessions and the 3, 1-hour punishment sessions. **F** Mean of the number of active and inactive nosepokes during the first hour of the last 3 acquisition sessions and the 3 punishment sessions. **G** Cocaine intake before treatment administration. **H** Number of infusions during the 3 post-punishment sessions and the cPNS session. **I** Number of active and inactive nosepokes during the 3 post-punishment sessions and the PS+ session. **J** Number of active nosepokes during the last post-punishment session and the PS+ session. **K** Intake during post-punishment self-administration. **L** Intake (Student’s *t* test) and **M** nosepokes during PS+ (ANOVA). **N** Latency to nosepoke in the active hole after the presentation of PS (Long-rank test). **O** Heatmap comparing resistant and sensitive groups. **P** Number of resistant and sensitive mice in each treatment group (Fisher’s exact test) (VEH n=21, URB n=20). Data are shown as mean ± SEM. ACQ: acquisition of cocaine self-administration, PNS: punishment sessions, SA: self-administration, PS+: conditioned punishment session, TO: time-out, VEH: vehicle, URB: URB597.

Subsequently, mice were divided in 2 groups that received either vehicle or URB597 (1 mg/kg) during three regular self-administration days (SA14-16) and one conditioned punishment session (PS+), where the predictive light was presented in the same way as in the punishment sessions, but no foot shock was delivered (Fig. 2A). A vehicle/URB597 injection was administered 30 minutes before each session (SA14-16 and PS+). Since punishment and PS+ sessions were limited to one hour, we considered only the first hour of the self-administration sessions for the purpose of comparisons. After the PS+ session, mice were euthanised and brains were dissected or perfused for further analyses.

#### Experiment 3: Assessment of URB597 modulation of drug-seeking behaviour during withdrawal

Mice were first trained to self-administer cocaine under FR1 (3-10 days) and FR3 (5 days) schedules of reinforcement. Mice progressed to a FR3 schedule when the criteria previously mentioned were met. Those who failed to meet the requirements within 10 days were excluded from the experiment.

Subsequently, mice were divided in 3 groups: withdrawal day 1 (WD1), WD30-VEH and WD30-URB. The WD1 group was tested for cue-induced drug-seeking behaviour the first day of withdrawal. Alternatively, the remaining two groups underwent a 30-day withdrawal in their home cages and were treated daily with either vehicle or URB597 for the first 25 days. Since we were interested in the URB597-mediated modulation of neuroplastic mechanisms underlying the expression of cocaine craving, a 5-day wash-out period prevented a potential bias of the acute effects of the treatment. On WD30, both groups were tested for cue-induced drug-seeking behaviour (Fig. 4A). During this 2-hour test, an active nosepoke led to the presentation of the contingent nosepoke cue, but no cocaine infusion was delivered. Only the first hour of the session was considered for the analyses to avoid the intrasession extinction phenomenon. Immediately after the seeking test, mice were dissected or perfused for further analyses.

### RT-qPCR

Total RNA extraction from ventral striatum (vSTR) samples was conducted using the trizol method as previously described [59,60]. For the qPCR, we used 20 ng of sample to evaluate the expression of CB1R, FAAH, mGluR5, NAPE-PLD and GAPDH as a housekeeping gene (see Supplementary material).

### Immunohistochemistry

This procedure was conducted as previously described with minor modifications (see Supplementary material) [61]. Free floating coronal sections of vSTR were incubated with primary antibodies against cFos, a neuronal activation marker, and CTIP2, a marker of MSN in striatum. For the labelling of the different interneurons, primary antibodies against parvalbumin (PV), somatostatin (STT) and choline acetyl transferase (ChAT) were used. The slices were subsequently incubated with the appropriate secondary antibodies (see supplementary Table 1 and 2). Images were obtained using sequential laser scanning confocal microscopy (Leica TCS SP5 upright) and quantified using Fiji-ImageJ software.

### Data and statistical analysis

Data are presented as mean ± SEM. We used GraphPad Prism 9.5.0 and IBM SPSS Statistics 29.0 software for statistical analysis and graphing. For analysis involving one, two or three factors, we used one-, two- or three-way analysis of variance (ANOVA), respectively. When an experimental condition followed a within-subject design, repeated measures ANOVAs were used. The factors considered were *treatment* (Vehicle/URB597), *group* (WD1/WD30-Vehicle/WD30-URB597), *time* (days or minutes), *session* (SA16/PS+) and *hole* (active/inactive). The statistical threshold for significance was set at p<0.05. When F achieved significance and there was no significant variance in homogeneity, a Tukey’s *post hoc* test was run. We analysed the results of single-factor experiments with two levels using unpaired Student’s *t* tests. Latency to nosepoke was analysed using Long-rank tests. Lastly, we used k-means cluster analyses on the entire population to identify resistant and sensitive mice to punishment (see Supplementary material). The number of resistant and sensitive mice in both the vehicle and URB597 groups was analysed using Fisher’s exact test.

## RESULTS

### URB597 does not modulate acquisition of cocaine self-administration

Administration of URB597 (1 mg/kg i.p.) 30 minutes before every acquisition session did not modulate cocaine seeking or taking (Fig. 1B-C).

### URB597 boosts conditioned punishment during cocaine self-administration

After acquisition of cocaine self-administration, mice were exposed to 3 daily punishment sessions, in which one third of the reinforcers were accompanied by the delivery of a foot shock and preceded by a predictive light (Fig. 2A-D). The comparison between the mean of infusions taken during the last three days of acquisition and those during the three days of punishment showed a dramatic punishment-induced reduction in drug seeking and taking (Fig. 2E-F).

To determine whether URB597 may modulate cocaine self-administration under conditioned punishment, mice were randomly assigned to the vehicle or URB597-group and underwent 3 days of post-punishment self-administration and a PS+ session. Cocaine consumption before treatment was not different between groups (Fig. 2G). Globally, the number of infusions and nosepokes during the sessions in which the treatments were administered was not different between groups (Fig. 2H-I). However, the comparison between SA16 and PS+ sessions revealed that both groups behaved differently on these days (Fig. 2J; two-way ANOVA: interaction *day* x *treatment* F_(1,39)_=4.185 p=0.047). In fact, cocaine intake during the post-punishment self-administration was not affected by the URB597 treatment (Fig. 2K). However, intake (Fig. 2L; Student’s *t* test: t_(39)_=1.849, p=0.072), but especially the number of nosepokes (Fig. 2M; two-way ANOVA: *hole* F_(1,39)_=26.81 p<0.001, *treatment* F_(1,39)_=6.136 p=0.017), were blunted in the URB597-treated during conditioned punishment. Specifically, only 30% of the URB597-treated mice responded after the first punishment-predictive stimulus of the session appeared, while 76% of the vehicle-treated mice did (Fig. 2N; Long-rank test: χ_2_=7.367 p=0.006). Finally, we used the k-means cluster analysis for the entire population to identify resistant and sensitive mice to punishment (Fig. 2O). These two populations differ in all the behavioural variables used for the cluster analyses, except inactive nosepokes. Importantly, the number of sensitive mice in the URB597-treated group was significantly higher compared to the vehicle-treated group (Fig. 2P; Fisher’s exact test p=0.012).

### URB597 increases CB1R gene expression and MSN activation in NAc shell

To test whether URB597 may affect molecular components that participate in the endocannabinoid-mediated plasticity in the vSTR, we assessed their gene expression and observed that URB597 increased CB1R gene expression (Fig. 3A; Student’s *t* test: t_(17)_=2.356, p=0.030).

**Fig 3.**
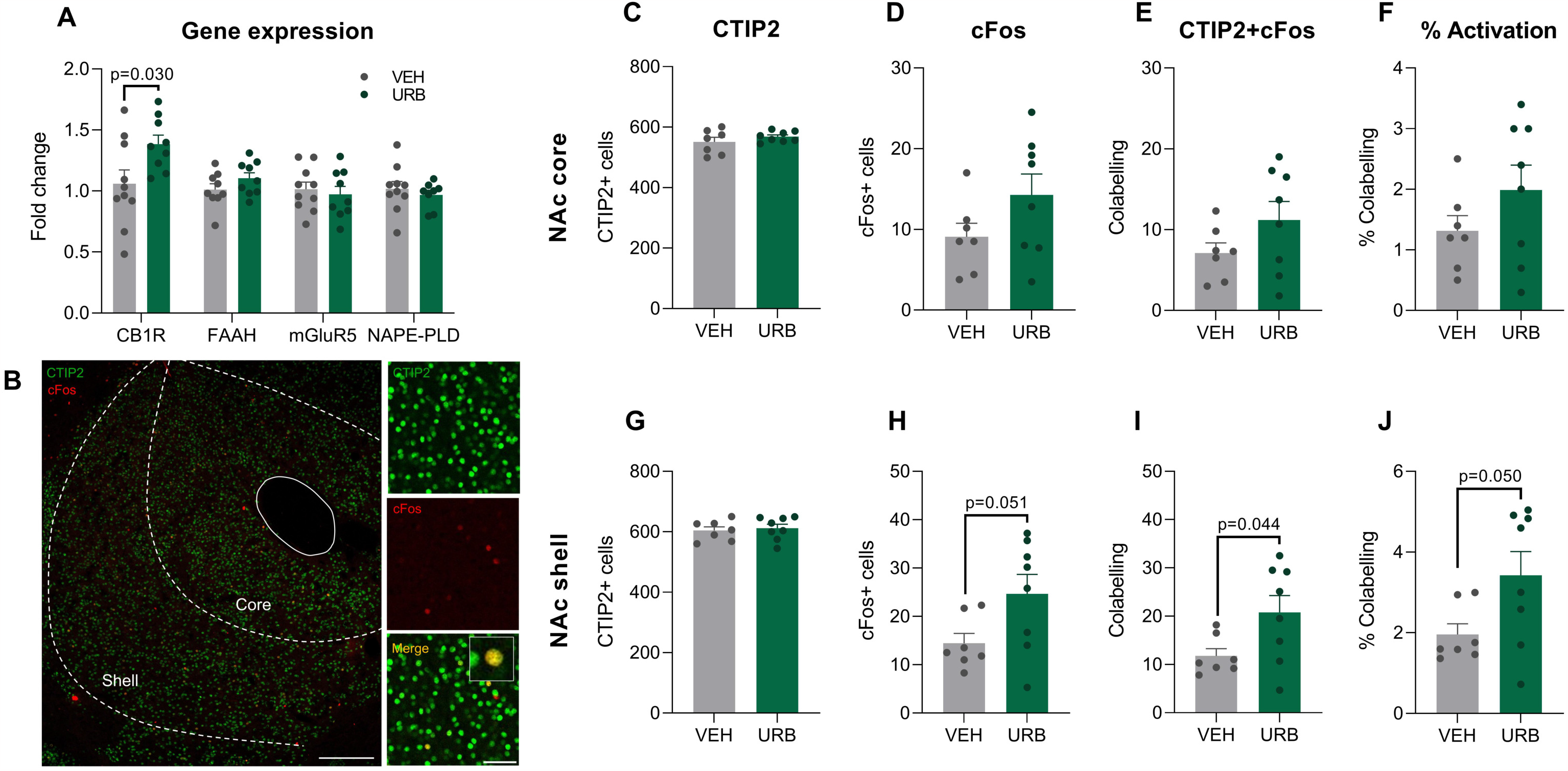
URB597 (1 mg/kg)-treated mice show an increase in MSN activation in NAc shell and CB1R gene expression in vSTR after cPNS. **A** Gene expression of CB1R, FAAH, mGluR5 and NAPE-PDL (Student’s *t* test) (n=8-10/group). **B** Confocal sections of NAc showing immunofluorescence for CTIP2 and cFos. Scale bar: 200 µm (left) and 50 µm (right). **C** Number of CTIP2+ and **D** cFos+ cells, **E** colocalization and **F** percentage of CTIP2-positive cells expressing cFos in NAc core and **G-J** NAc shell (Student’s *t* test) (n=7-8/group). Data are shown as mean ± SEM. VEH: vehicle, URB: URB597, CB1R: cannabinoid receptor 1, FAAH: fatty acid amide hydrolase, mGluR5: metabotropic glutamate receptor 5, NAPE-PLD: N-acyl phosphatidylethanolamine-specific phospholipase D; CTIP2: transcription factor COUP TF1-interacting protein 2.

**Fig 4.**
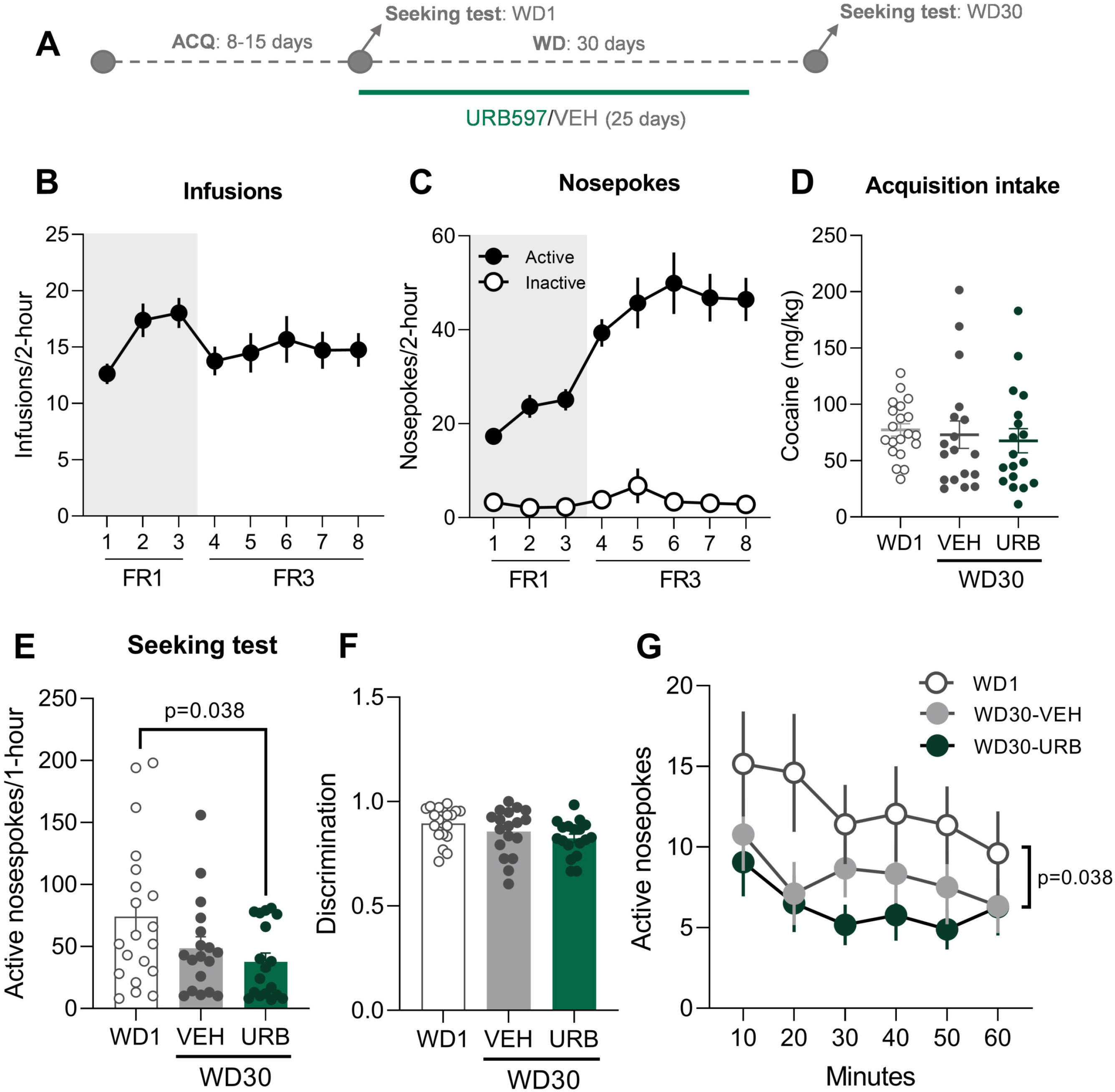
URB597 (1 mg/kg) decreases cocaine seeking during abstinence from cocaine self-administration. **A** Schematic timeline of the experiment 3. **B** Infusions and **C** nosepokes performed during the acquisition of cocaine self-administration (n=56). **D** Cocaine intake before treatment administration. **E** Active nosepokes (Tukey) and **F** discrimination during the cue-induced drug-seeking tests. **G** Active nosepokes (Tukey) and throughout the first hour of the seeking test (Tukey) (WD1 n=20, WD30-VEH n=18, WD30-URB n=18). Data are shown as mean ± SEM. ACQ: acquisition of cocaine self-administration, WD: withdrawal day, VEH: vehicle, URB: URB597, FR: fixed ratio.

Besides, we quantified the number of cells expressing CTIP2 and cFos in the NAc after the PS+ session (Fig. 3B). In the NAc shell, we found an increase in cFos expression due to URB597 treatment located in the MSNs (Fig. 3G-J; Student’s *t* test: cFos-positive (+) cells t_(13)_=2.147, p=0.051, colocalization t_(13)_=2.218, p=0.044, % activation t_(13)_=2.160, p=0.050), but there were no significant differences in the NAc core (Fig. 3C-F).

### URB597 treatment mildly mitigates drug-seeking behaviour during cocaine withdrawal

To further characterize URB597 effects on cocaine-induced addictive behaviours, we treated mice with either vehicle or URB597 during the withdrawal period and tested them for cue-induced drug-seeking behaviour (Fig. 4A). Mice were trained to self-administer cocaine under FR1 and FR3 to promote drug-seeking behaviour (Fig. 4B-C). Cocaine consumption prior to treatment was not different among groups (Fig. 4D). The results obtained in the drug-seeking tests revealed that the groups behaved differently (Fig. 4E; one-way ANOVA: F_(2,53)_=3.386 p=0,041). The URB597-treated group significantly decreased drug seeking compared to WD1 group (Tukey: p=0.038), whereas the VEH-treated group did not show changes. Similarly, the discrimination index (active nosepokes/total nosepokes) followed the same trend, however, the differences found did not reach statistical significance (Fig. 4F; one-way ANOVA: *group* F_(2,53)_=2.769 p=0,071).

The active nosepokes registered throughout the first 60 minutes of the test revealed that the three groups behaved differently (Fig. 4G; two-way ANOVA: *time* F_(4.260,225.8)_=2.368 p=0.049, *group* F_(2,53)_=3.386 p=0.041). Specifically, the *post hoc* for *group* showed that the URB597-treated mice had a lower response rate compared to the WD1 group (Tukey, p=0.038), suggesting that URB597-treated mice displayed a lower craving expression severity.

### URB597 administered during withdrawal prevents the increase in FAAH gene expression and reduces MSNs activation in the NAc

We evaluated the expression of different genes in the vSTR and we found that FAAH expression was increased after 30 days of withdrawal (Fig. 5A; one-way ANOVA: F_(2,19)_=3.526 p=0,049, Tukey WD1 vs WD30-VEH p=0.045), an effect that was prevented by the administration of URB597.

**Fig 5.**
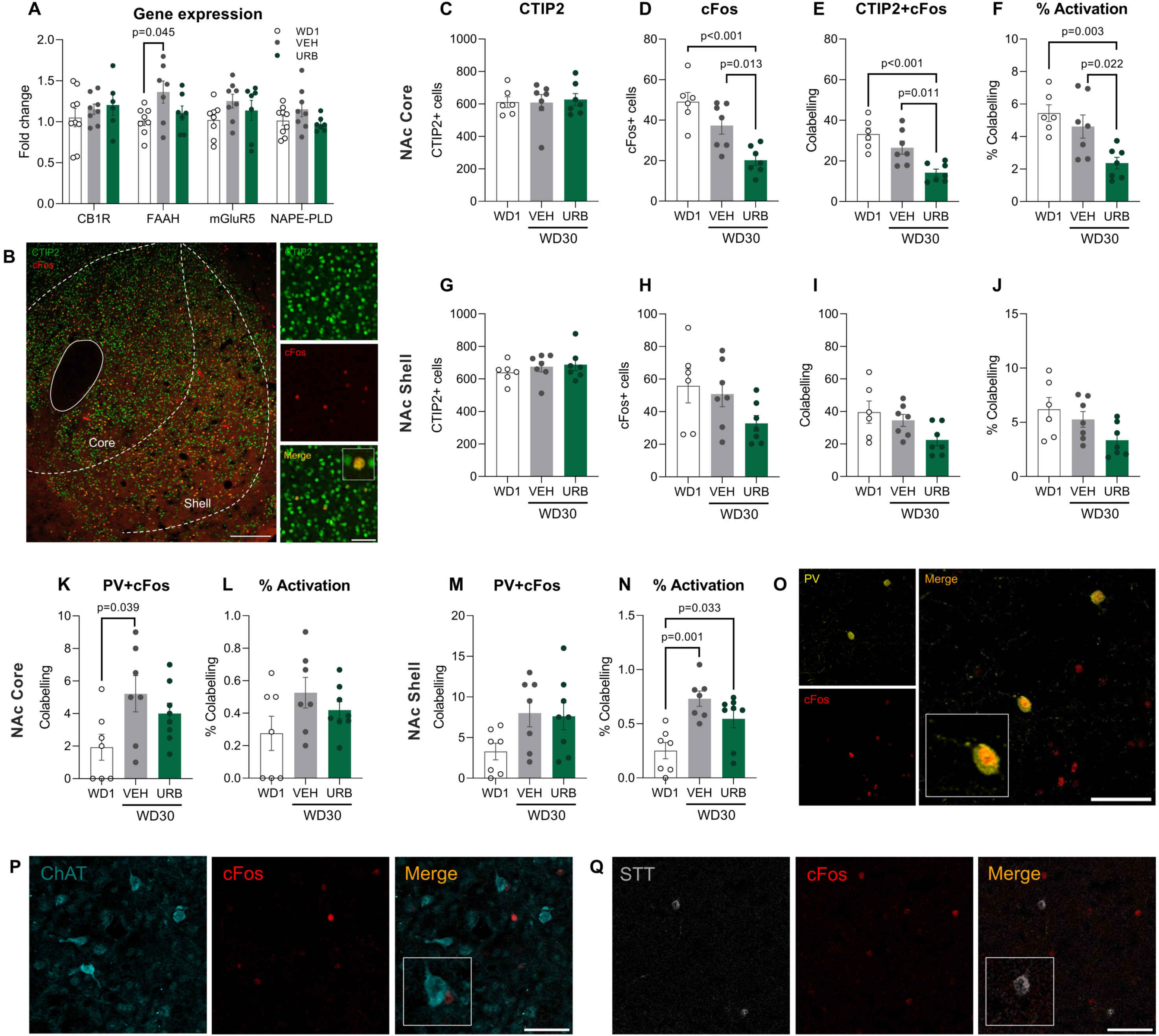
URB597 (1 mg/kg) administered during withdrawal blunts MSN activation in NAc and prevents the increase of FAAH in vSTR. **A** Gene expression of CB1R, FAAH, mGluR5 and NAPE-PDL (Tukey) (n=6-9/group). **B** Confocal sections of NAc showing immunofluorescence for CTIP2 and cFos. Scale bar: 200 µm (left) and 50 µm (right). **C** Number of CTIP2+ and **D** cFos+ cells, **E** colocalization and **F** percentage of CTIP2-positive cells expressing cFos in NAc core (Tukey) and **G-J** NAc shell (Tukey) (n=6-7/group). **K** Number of cells expressing PV and cFos (Tukey) and **L** the percentage of PV+ cells expressing cFos in NAc core and **M-N** NAc shell (Tukey) (n=6-7/group). **O** Confocal sections of NAc showing immunofluorescence for PV and cFos. Scale bar: 50 µm. **P** Confocal sections of NAc showing immunofluorescence for ChAT or **Q** STT and cFos. Scale bar: 50 µm. Data are shown as mean ± SEM. VEH: vehicle, URB: URB597, CB1R: cannabinoid receptor 1, FAAH: fatty acid amide hydrolase, mGluR5: metabotropic glutamate receptor 5, NAPE-PLD: N-acyl phosphatidylethanolamine-specific phospholipase D; CTIP2: transcription factor COUP TF1-interacting protein 2, PV: parvalbumin, ChAT: choline acetyl transferase, STT: somatostatin.

NAc core immunofluorescence images showing CTIP2+ and cFos+ cells (Fig. 5B) revealed a decrease in the number of cFos+ cells (Fig. 5D; one-way ANOVA: F_(2,17)_=14 p<0.001; Tukey WD1 vs WD30-VEH p<0.001 and WD30-VEH vs WD30-URB p=0.013), a decrease in the colocalization with CTIP2 (Fig. 5E; one-way ANOVA: F_(2,17)_=12.33 p<0.001; Tukey WD1 vs WD30-VEH p<0.001 and WD30-VEH vs WD30-URB p=0.011) and in the percentage of MSN co-expressing cFos (Fig. 5F; one-way ANOVA: F_(2,17)_=8.351 p=0.003; Tukey WD1 vs WD30-VEH p=0.003 and WD30-VEH vs WD30-URB p=0.022) in the URB597-treated group compared to both WD1 and WD30-VEH. This effect was also observed in the NAc shell, although less pronounced and was not statistically significant (Fig. 5H-J; one-way ANOVA: F_(2,17)_=3.355 p=0.059).

For a better understanding of the URB597 effects on neuronal activation, we assessed cFos expression in three types of interneurons (PV+, STT+ and ChAT+). In the NAc core, we found an increase in the number of neurons expressing both PV and cFos after 30 days of withdrawal that was prevented by URB597 (Fig. 5K; one-way ANOVA: F_(2,19)_=3.619 p=0.046, Tukey WD1 vs WD30-VEH p=0.039). However, the percentage of PV+ cells expressing cFos was increased in both WD30-VEH and WD30-URB compared to WD1 in the NAc shell (Fig. 5N-O; one-way ANOVA: F_(2,19)_=9.54 p=0.001, Tukey WD1 vs WD30-VEH p=0.001, WD1vs WD30-URB p=0.033), but not the NAc core (Fig. 5L). No colocalization of ChAT or STT with cFos was found (Fig. 5P-Q).

## DISCUSSION

The present study provides evidence of the modulation caused by FAAH inhibition on cocaine reinforcement. URB597 treatment mitigates both cocaine consumption under the risk of punishment and cue-induced drug-seeking behaviour during protracted abstinence, while having no impact on acquisition of cocaine self-administration. These behavioural outcomes are accompanied by changes in MSNs activation in the NAc as well as CB1R and FAAH gene expression in the vSTR.

In agreement with previous studies that administered URB597 acutely (0.1-3 mg/kg) [53,54], our results indicate that a repeated URB597 (1 mg/kg) treatment during the acquisition of cocaine self-administration does not modulate drug taking (Fig. 1). To date, URB597 treatment (acute, 0.5 mg/kg) has been shown to decrease alcohol intake in a mouse model of intermittent access [47]. Otherwise, it failed to alter tetrahydrocannabinol (THC) [54] and cocaine [53,54] self-administration as well as sucrose intake [47], and it increased AEA self-administration [54]. These discrepancies indicate that URB597 effects are reinforcer-dependent. However, the congruency of our data with previous studies strongly suggests that URB597 does not interfere with the acquisition of cocaine self-administration.

To examine the effects of FAAH inhibition on compulsive drug taking, measured cocaine intake under conditioned punishment. In this experiment (Fig. 2), URB597 was administered during post-punishment self-administration and a PS+ session. This strategy has two advantages: (i) it offers a translational approach because URB597 treatment is applied after the consolidation of drug taking and the appearance of the associated negative consequences, and (ii) considering the analgesic and anxiolytic effects of URB597 [23–25], we avoid biases in the interpretation of the results by circumventing the contingency between treatment and punishment.

While URB957 failed to affect cocaine self-administration after punishment, it did decrease cocaine consumption upon presentation of the punishment-predictive stimulus, suggesting that URB597 treatment enhances sensitivity to punishment. Accordingly, most of the mice treated with URB597 displayed a sensitive phenotype to punishment and, therefore, a robust pavlovian conditioned inhibition of drug intake. In contrast, the predictive stimulus failed to elicit such behaviours in most of the control mice. The mechanism by which FAAH inhibition facilitates the supressing effects of the punishment-predictive stimulus in the context of cocaine self-administration remains unclear. We hypothesize that this could be related to a facilitation of the aversive memory retrieval. Consistent with this hypothesis, a recent study found that local blockade of TRPV1 in CA1 of hippocampus prevented the retrieval of contextual fear memories (measured by freezing behaviour) of mice exposed to foot shocks and reported a positive correlation between the levels of AEA and the freezing behaviour [62]. The authors propose opposite roles of CB1R and TRPV1 in the modulation of conditioned fear [63,64], in which activation of TRPV1 by AEA would lead to the strengthening of aversive memories. Another possible explanation could be related to a URB597-induced improvement in properly assigning value to aversive outcomes that occur as a consequence of compulsive drug seeking [65].

Additionally, URB597-treated mice exhibited an increase of CB1R gene expression in the vSTR. Similarly, we previously reported that cannabidiol (20 mg/kg), which acts as FAAH inhibitor, when administered in the context of cocaine self-administration, increases CB1R protein levels in vSTR and in other brain regions involved in cocaine use [66,67]. Certainly, additional research would be needed to determine whether the increase in conditioned suppression of cocaine intake following URB597 administration is mediated by CB1R. Besides, we found a higher activation of MSNs in the NAc shell of URB597-treated mice. The NAc shell is strongly involved in the control of punishment-induced suppression of reward seeking [14,15]. NAc shell inactivation invigorates punished responding for food [15], suggesting that it mediates inhibition of punished reward seeking. Glutamatergic projections from mPFC [68,69] and amygdala [15,70,71], especially the caudal basolateral amygdala (BLA) [72], to the NAc shell contribute to punishment avoidance, adjusting behaviour in response to consequences. Importantly, URB597 (0.1 mg/kg) prevented the cocaine-induced inhibition of MSNs excitability in the BLA-NAc shell pathway [44], further supporting its facilitating effects in the MSNs activation under risk of punishment. Our results, together with previous literature, strengthen the therapeutical potential of FAAH inhibition for the treatment of compulsive-like cocaine intake.

In a similar line, we also demonstrated that FAAH inhibition has a role in the expression of craving during cocaine abstinence (Fig. 4). In accordance, URB597 administered acutely after extinction training [53] or chronically during forced abstinence [55], decreased cue-induced seeking behaviour in rats. However, the URB597-induced inhibitory effects were subtle in our experimental conditions. Although the URB597-treated group exhibited a noteworthy reduction in drug seeking during withdrawal, it did not differ significantly with the vehicle group, indicating that the treatment’s impact is relatively modest.

The reduction in cocaine-seeking behaviour observed with URB597 treatment was linked to changes in FAAH gene expression and neuronal activation in the NAc. Since the ECS is intricately involved in reward processing and motivation [5,7], exposure to and withdrawal from cocaine self-administration can profoundly impact its functioning. For example, studies in rats have shown that, after cocaine self-administration, AEA levels in the NAc are reduced [73], even after 10 days of extinction training [74]. Consistent with these findings, we observed a withdrawal-induced increment in FAAH gene expression, which would eventually lead to decreased AEA levels due to its increased degradation. Interestingly, URB597 treatment during abstinence blocked the increase in FAAH gene expression. Likewise, electrophysiological studies have shown that exposure to cocaine impairs eCB-LTD in the glutamatergic synapses within the NAc [9,10,75], even after periods of withdrawal [8,76]. Hence, Grueter et al. (2010) demonstrated that this type of plasticity is mediated by AEA through two complementary mechanisms: (i) activation of CB1R in the presynaptic neuron, which leads to decreased glutamate release, and (ii) activation of TRPV1 in the postsynaptic neuron, which causes endocytosis of α-amino-3-hydroxy-5-methyl-4-isoxazolepropionic acid receptors (AMPARs) [10]. Notably, URB597 facilitates both types of plasticity, resulting in an increased eCB-LTD in the NAc core [10]. In fact, direct activation of TRPV1 by capsaicin can prevent the cocaine-induced loss of plasticity. This evidence, together with our results that suggest a blunted degradation of AEA due to URB597 treatment during withdrawal, have led us to hypothesize that URB597 might reduce MSNs activation in the NAc core by restoring eCB-LTD. A large body of literature supports the prominent role of NAc core in driving approach behaviours [11,13], including those related to motivationally-relevant stimuli such as drug-associated cues [16]. Accordingly, we found that a URB597-induced decrease in MSNs activation in this brain region is associated with a mitigation of cue-induced cocaine-seeking behaviour.

To further understand the effects of URB597 in NAc activation during the cue-induced drug seeking test, we assessed cFos expression in the NAc interneurons, which make up the 5% of the neuronal population. They are mainly classified as (i) fast-spiking parvalbumin-releasing (PV+), (ii) persistent low threshold somatostatin-releasing (STT+), and (iii) tonically active acetylcholine-releasing (ChAT+) interneurons [77,78]. These interneurons work in a complementary way to control the activity of MSNs and there is evidence of their implication in cocaine-related behaviours [79–84]. Accordingly, we found an increase in the activation of PV+ interneurons after 30 days of cocaine abstinence. Because PV+ interneurons exert inhibitory control over MSNs [85], an increase in their activation could be related to the reduced MSNs activity observed. However, although 80% of PV+ interneurons express CB1R [86] and their synapses onto MSNs are regulated by eCB-LTD [85], URB597 did not exert any effect over their activation. This indicates that its actions on cue-induced cocaine seeking during abstinence are not mediated by the modulation of PV+ interneurons. Finally, STT+ and ChAT+ interneurons showed no activation.

In conclusion, we have demonstrated a previously unknown role of FAAH inhibition in compulsive cocaine taking and extended the evidence about its anti-relapse effects. Interestingly, the FAAH inhibitor URB597 does not appear to modulate the primary reinforcing effects of cocaine *per se*, but instead it reduces drug-seeking behaviour when a negative component comes into play (conditioned punishment and withdrawal). These findings have allowed us to propose that URB597 might exert its anti-addictive effects by strengthening the sensitization to negative outcomes associated to cocaine use. Lastly, our research supports the potential of FAAH inhibition as a promising therapeutic target for the treatment of cocaine use disorder.

## ACKNOWLEDGEMENTS

We would like to thank Xavier Puig-Reyné for his technical assistance during the experimental procedures.

## AUTHOR CONTRIBUTION

L.A.-Z. and O.V. were responsible for the study, concept, and design. L.A.-Z., A.G.-B, A.C.-Z, I.G.-L. and A.M.-S. carried out the behavioural procedures. L.A.-Z. carried out the molecular and histological procedures L.A.-Z. and O.V. drafted the manuscript. All authors critically reviewed the content and approved the final version for publication.

## FUNDING

This work was supported by the Ministerio de Ciencia e Innovación (#PID2019-104077RB-100 MCIN/AEI/10.13039/501100011033), Ministerio de Sanidad (Plan Nacional sobre Drogas #2018/007 and ISCIII-Feder-RIAPAd-RICORS #RD21/0009/001) and by the Generalitat de Catalunya, AGAUR (#2021SGR00485). L.A-Z received a FPI grant (BES-2017-080066) associated to #SAF2016-75966-R grant to O.V. from Ministerio de Economia y Competitividad. A.G-B received a FI-AGAUR grant from the Generalitat de Catalunya (2019FI_B0081). I.G-L. obtained a grant from the Ministerio de Ciencia e Innovación (PRE2020-091923). The Department of Medicine and Health Sciences (UPF) is a “Unidad de Excelencia María de Maeztu” funded by the AEI (#CEX2018-000792-M). O.V. is recipient of an ICREA Academia Award (Institució Catalana de Recerca i Estudis Avançats, Generalitat de Catalunya).

## COMPETING INTERESTS

The authors have nothing to disclose.

## SUPLEMENTARY MATERIAL

### Surgery

Surgical implantation of the catheter into the jugular vein was performed following anaesthetization with a mixture of ketamine hydrochloride (75 mg/kg; Imalgène1000, Lyon, France) and medetomidine hydrochloride (1 mg/kg; Medeson®, Barcelona, Spain). The anaesthetic solution was injected in a volume of 0.15 mL/10 g body weight, i.p. Briefly, a 6 cm length of silastic tubing (0.3 mm inner diameter, 0.6 mm outer diameter) (silastic, Dow Corning, Houdeng-Goegnies, Belgium) was fitted to a 22-gauge steel cannula (Semat, Herts, England) that was bent at a right angle and then embedded in a cement disk (Dentalon Plus, Heraeus Kulzer, Germany) with an underlying nylon mesh. The catheter tubing was inserted 1.3 cm into the right jugular vein and anchored with a suture. The remaining tubing ran subcutaneously to the cannula, which exited at the mid-scapular region.

Meloxicam (0.5 mg/kg s.c.; Metacam®, Barcelona, Spain), enrofloxacin (7.5 mg/kg i.p.; Baytril® 2.5%; Barcelona, Spain), atipamezole hydrochloride (0.5 mg/kg i.p.; Revertor®, Barcelona, Spain), and 1 ml glucose 5% solution were injected after the surgery. Home cages were placed on thermal blankets to avoid post-anaesthesia hypothermia. Mice were daily monitored for their weight and treated with meloxicam for 48 h. Mice were allowed to recover for at least four days before the acquisition phase of the self-administration procedure began.

### Acquisition of cocaine self-administration

The self-administration experiments took place in operant conditioning chambers (Model ENV-307A-CT, Med Associates, Inc. Cibertec. Madrid. Spain) with two nosepoke holes. Animals were connected to a microinfusion pump (PHM-100A, Med-Associates, Georgia, VT, USA). Active and inactive holes were selected randomly for each operant box. A nosepoke in the active hole resulted in the delivery of a 20 μl cocaine infusion over 2 seconds (0.6 mg/kg/inf) and a stimulus light for 4 seconds. Nosepokes in the inactive hole had no consequences but were registered for the analyses. Each infusion was followed by a 15-second time-out period. All sessions started with a priming infusion and lasted 2 hours.

### RT-qPCR

Total RNA extraction from vSTR samples was conducted using trizol. rt-PCR was performed by High-Capacity cDNA Reverse Transcription Kit (Applied Biosystems, Foster City, CA) using random primers and following standardized protocols. As per the qPCR, 20 ng of sample were loaded in addition to the following reagents: LightCycler SBYR green 480 Master Mix (Roche LifeScience, Product No. 04707516001) and the specific primers for the target genes (Integrated DNA Technologies, Inc.). The qPCR was performed in LightCycler® 480 Instrument II (Roche LifeScience).

### Immunohistochemistry

Free-floating sections were washed three times for 5 min with 0.05 M TRIS buffered saline pH 7.6 (TBS). In brief, sections were: (i) pre-incubated in 4% normal goat/donkey serum in 0.05 M TBS pH 7.6 with 1% Triton X-100, at RT for 1 h, to block nonspecific labelling; (ii) incubated in primary antibodies (see Supplementary Table 1) diluted in 0.05 M TBS at 4°C for 24 h; (iii) incubated with fluorescent-labelled secondary antibodies (see Supplementary Table 2) diluted in 0.05 M TBS for 2 h at RT. To reveal the cytoarchitecture, sections were counterstained prior to mounting by bathing them for 5 min in Hoechst (a nuclear staining, 1:10000; Invitrogen, Cat#33258) at RT. After each step, sections were washed three times for 5 min in 0.05 M TBS except between steps (i) and (ii). Finally, sections were washed in 0.05 M TB, mounted onto gelatinised slides and cover-slipped with fluorescence mounting medium (FluorSave Reagent, Merck Milipore, Cat#345789, Darmstadt, Germany). Images were obtained using sequential laser scanning confocal microscopy (Leica TCS SP5 upright) and a 20x objective. Photographs were taken in both hemispheres. The number of cells expressing each marker and the colocalization with cFos was quantified using Fiji-ImageJ software. Percentage of activation was calculated as follows: (number of colocalized cells/number of CTIP2+/PV+ cells) x 100.

**Supplementary Table 1.** Primary antibodies used for immunohistochemistry.

| Antobody | Description | Host | Dilution | Company | Item number |
| --- | --- | --- | --- | --- | --- |
| cFos | cFos | Mouse | 1:1000 | Abcam | ab208942 |
| CTIP2 | Transcription factor COUP<br>TF1-interacting protein 2 | Rabbit | 1:500 | Atlas Antibodies | HPA049117 |
| PV | Parvalbumin | Guinea Pig | 1:1000 | Synaptic Systems | 195 004 |
| ChAT | Choline acetyl transferase | Goat | 1:500 | Merck Millipore | AB144 |
| STT | Somatostatin | Rat | 1:500 | Merck Millipore | MAB354 |

**Supplementary Table 2.**
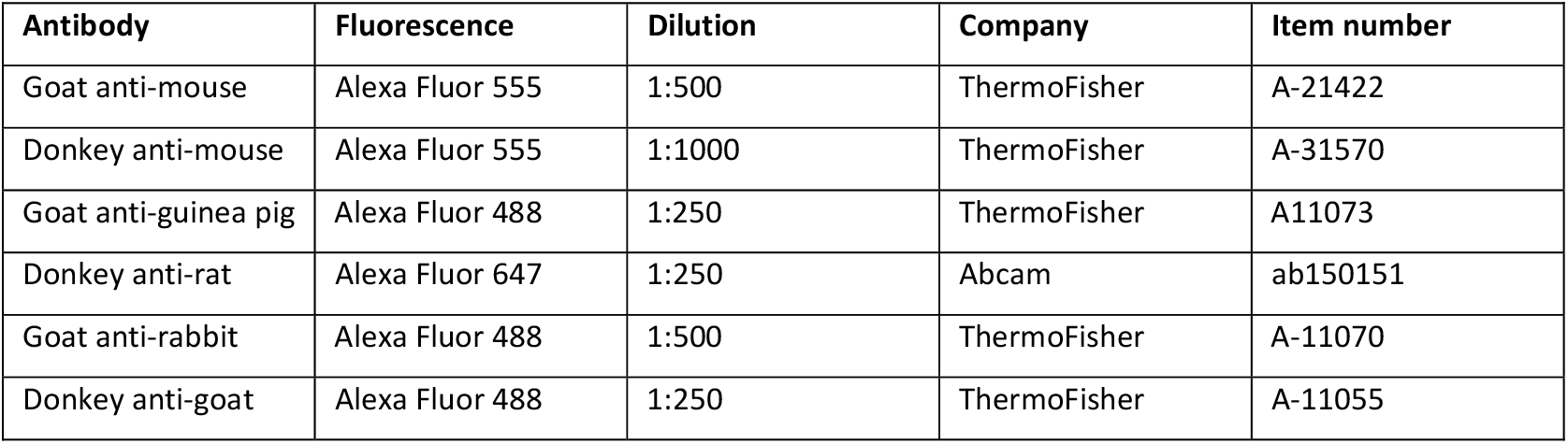
Secondary antibodies used for immunohistochemistry.

| Antibody | Fluorescence | Dilution | Company | Item number |
| --- | --- | --- | --- | --- |
| Goat anti-mouse | Alexa Fluor 555 | 1:500 | ThermoFisher | A-21422 |
| Donkey anti-mouse | Alexa Fluor 555 | 1:1000 | ThermoFisher | A-31570 |
| Goat anti-guinea pig | Alexa Fluor 488 | 1:250 | ThermoFisher | A11073 |
| Donkey anti-rat | Alexa Fluor 647 | 1:250 | Abcam | ab150151 |
| Goat anti-rabbit | Alexa Fluor 488 | 1:500 | ThermoFisher | A-11070 |
| Donkey anti-goat | Alexa Fluor 488 | 1:250 | ThermoFisher | A-11055 |

### Statistical analyses

We used k-means cluster analyses on the entire population to identify resistant and sensitive mice to punishment. For this analysis we considered behavioural variables performed during the post-punishment self-administration (intake, active nosepokes, active nosepokes during time-out and inactive nosepokes) and PS+ session (intake, active nosepokes, inactive nosepokes and latency to nosepoke after the presentation of PS). The number of resistant and sensitive mice in the vehicle and URB597 groups were analysed using Fisher’s exact test.

**Supplementary Fig 1.**
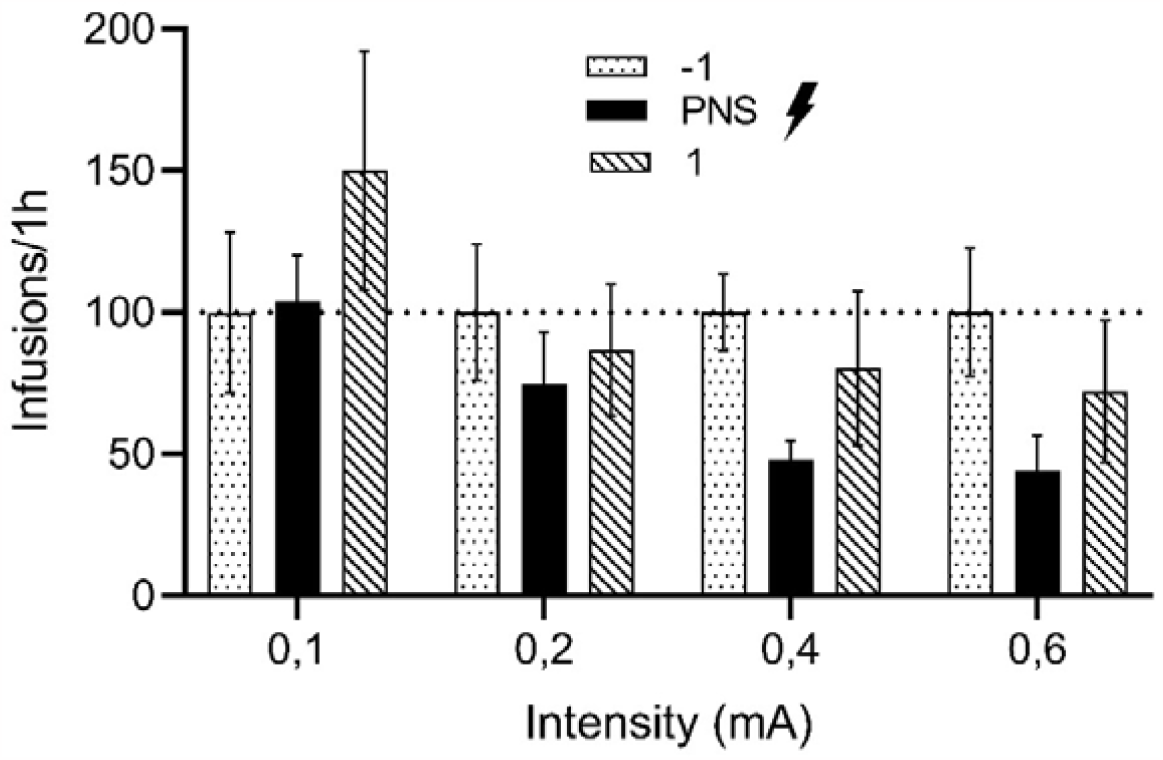
Cocaine consumption after application of electric foot shock. After acquisition of cocaine self-administration, different groups of mice (n=5-7/group) were exposed to different foot shock intensities (0.1, 0.2, 0.4 and 0.6 mA). We show the number of infusions taken during the punishment session (PNS), the self-administration day before (−1) and after (1). 0.4 mA was selected as the preferred intensity for the following experiments for standing as the lowest one that decreases cocaine consumption in more than 50%, and it maintains part of its inhibitory effects on the following self-administration day.

## Notes

### Competing Interest Statement

The authors have declared no competing interest.

